# Multiple performance peaks for scale-biting in an adaptive radiation of pupfishes

**DOI:** 10.1101/2023.12.22.573139

**Authors:** Anson Tan, Michelle St. John, Dylan Chau, Chloe Clair, HoWan Chan, Roi Holzman, Christopher H. Martin

**Author notes:** Corresponding author: Christopher Martin, 3101 Valley Life Sciences Building, University of California, Berkeley 94720.

## Abstract

The physical interactions between organisms and their environment ultimately shape their rate of speciation and adaptive radiation, but the contributions of biomechanics to evolutionary divergence are frequently overlooked. Here we investigated an adaptive radiation of *Cyprinodon* pupfishes to measure the relationship between feeding kinematics and performance during adaptation to a novel trophic niche, lepidophagy, in which a predator removes only the scales, mucus, and sometimes tissue from their prey using scraping and biting attacks. We used high-speed video to film scale-biting strikes on gelatin cubes by scale-eater, molluscivore, generalist, and hybrid pupfishes and subsequently measured the dimensions of each bite. We then trained the SLEAP machine-learning animal tracking model to measure kinematic landmarks and automatically scored over 100,000 frames from 227 recorded strikes. Scale-eaters exhibited increased peak gape and greater bite length; however, substantial within-individual kinematic variation resulted in poor discrimination of strikes by species or strike type. Nonetheless, a complex performance landscape with two distinct peaks best predicted gel-biting performance, corresponding to a significant nonlinear interaction between peak gape and peak jaw protrusion in which scale-eaters and their hybrids occupied a second performance peak requiring larger peak gape and greater jaw protrusion. A bite performance valley separating scale-eaters from other species may have contributed to their rapid evolution and is consistent with multiple estimates of a multi-peak fitness landscape in the wild. We thus present an efficient deep-learning automated pipeline for kinematic analyses of feeding strikes and a new biomechanical model for understanding the performance and rapid evolution of a rare trophic niche.

## Introduction

Most biodiversity evolved during adaptive radiation, the process by which a single lineage colonizes myriad ecological niches and rapidly diversifies (Gillespie et al., 2020; Martin and Richards, 2019; Simpson, 1944; Stroud and Losos, 2016). However, this process of rapid adaptation is also constrained by physical laws governing interactions between the organism and the essential tasks needed for survival and reproduction within its environment, known as the emerging field of ecomechanics (Higham et al., 2016; Higham et al., 2021; Perevolotsky et al., 2020). Understanding how functional traits interact in complex ways to achieve these essential life history tasks is fundamental for connecting population genomics to phenotypes to performance to fitness to adaptive radiation within new environments (Arnold, 1983; Martin et al., 2019). For example, estimating the performance landscape for a functional task or tasks can lead to deep insights about patterns of morphospace occupation and rates of morphological diversification across diverse taxa (Armbruster, 1990; Benkman, 1993; Figueirido et al., 2013; Holzman et al., 2022; Keren et al., 2018; Olsson et al., 2020; Raup, 1966; Schultz et al., 2021; Stayton, 2019; Tseng and Flynn, 2018). However, few systems have been investigated across these levels of biological organization.

Scale-biting, or lepidophagy, is a fascinating ecological niche that has evolved at least 20 times in fishes and within a diverse range of environments, from rift lakes (Hori, 1993; Raffini et al., 2017; Stewart and Albertson, 2010; Takahashi et al., 2007) to tropical streams and rivers (Evans et al., 2017; Gosavi et al., 2018; Gosavi et al., 2019; Grubh et al., 2004) to the mesopelagic zone (Nakae and Sasaki, 2002) and across ontogenetic stages (Davis et al., 2011; MacLeod, 2020; Novakowski et al., 2004; Szelistowski, 1989) in a diverse range of fish groups, including cichlidiformes, characiformes, siluriformes, and sharks (Kolmann et al., 2018; Martin and Wainwright, 2013c; Sazima, 1984). This trophic niche provides an excellent setting for investigating the functional traits underpinning performance and fitness because successful scale-eating appears to require both intensive prey capture behaviors and efficient energy use per strike (Sazima, 1984). In contrast to piscivory, all scale-eaters are generally smaller than their larger prey and numerous strikes must be performed efficiently due to the low energy payoff in calories per strike, resulting in an excellent laboratory and field model for observing repeated prey capture behaviors. Scale-eaters rarely or never consume whole prey (Gosavi et al., 2018; Kovac et al., 2019; Martin and Wainwright, 2013c; Sazima, 1984; Sazima, 1986).

The scale-biting pupfish, *Cyprinodon desquamator*, is the youngest lepidophage so far discovered; this species evolved within the past 10-15 kya on San Salvador Island, Bahamas (Martin and Wainwright, 2013a; Martin et al., 2019). Scales comprise approximately 50% of its diet in addition to macroalgae and microinvertebrates in several hypersaline lakes where it is endemic (Martin and Wainwright, 2011; Martin and Wainwright, 2013c; McLean and Lonzarich, 2017). All other extant scale-eating lineages likely evolved at least 1 Mya (Koblmüller et al., 2007), except for the extinct Lake Victorian lepidophagous cichlid *Haplochromis welcommei* (Greenwood, 1965). In several interior hypersaline lakes on San Salvador Island (SSI), *C. desquamator* occurs in sympatry at low frequencies in the same benthic macroalgae-dominated habitats as *C. variegatus*, a widespread generalist species, *C. brontotheroides*, an endemic oral-shelling molluscivore species, and *C.* sp. ‘wide-mouth’, a newly discovered intermediate scale-eating ecomorph which is not included in this study (Hernandez et al., 2018; Martin and Feinstein, 2014; Richards and Martin, 2022; St John et al., 2020).

The SSI adaptive radiation exhibits one of fastest rates of craniofacial diversification among any measured vertebrate group, up to 1,000 times faster than generalist populations on neighboring Bahamian islands for oral jaw length, due to rapid adaptation to the novel trophic niche of lepidophagy (Martin, 2016b; Martin and Wainwright, 2011). Previous field experiments measuring the growth and survival of F2 and F5 hybrids among all three species placed in field enclosures within two hypersaline lakes on San Salvador Island estimated a two-peak fitness landscape and a large fitness valley isolating the hybrids with greatest phenotypic similarity to scale-eaters (Martin and Wainwright, 2013b; Patton et al., 2022). This two-peak landscape was stable over multiple years, lake environments, and phenotype-frequency manipulations, suggesting that biophysical constraints on the interaction between pupfish craniofacial traits and their bite performance on scales may ultimately shape the adaptive landscape in this system, rather than frequency-dependent competition (e.g. (Hori, 1993; Martin, 2012)) or seasonal resource abundance (Grant and Grant, 2002), which would predict a dynamically changing landscape (Martin, 2016a; Martin and Gould, 2020). Furthermore, our genomic and developmental genetic analyses suggest that introgression of genetic variation for traits related to feeding behavior triggered adaptive radiation on SSI and led to the reassembly of Caribbean-wide genetic variation into a specialized scale-eater and molluscivore on a single island over several thousand years (McGirr and Martin, 2017; McGirr and Martin, 2018; Palominos et al., 2023; Richards and Martin, 2017; Richards et al., 2021).

In our initial pilot study of scale-biting kinematics in this system, we found that scale-eaters behaviorally reduced their realized peak gape during feeding strikes by reducing their jaw opening angle (St. John et al., 2020). However, we were unable to robustly quantify the performance landscape with limited strike and landmark data and did not measure F2 hybrids. Here we used all new feeding videos filmed at substantially improved resolution and developed a new automated machine-learning pipeline to quantify five landmarks on nearly every frame. We sampled extensive phenotypic diversity within the radiation using both purebred species and multiple F2 hybrid intercrosses and backcrosses to estimate the five-dimensional kinematic performance landscape for scale-biting.

## Methods

### Collection and Husbandry

Using seine nets or hand nets, we collected molluscivore (*C. brontotheroides*) and scale-eating (*C. desquamator*) pupfish from Crescent Pond, Little Lake, and Osprey Lake on San Salvador Island, Bahamas, and generalist (*C. variegatus*) pupfish from Lake Cunningham on New Providence Island, Bahamas and Fort Fisher estuary in North Carolina, United States in 2017 and 2018. Generalist pupfish from SSI could not be readily trained to feed on gelatin cubes during the period of this study; however, generalists from neighboring islands exhibit essentially identical craniofacial morphology and kinematics relative to SSI generalists (Martin, 2016b; St. John et al., 2020) . Wild-caught and lab-reared fish were maintained in 40-80 l aquaria at salinities of 2-8 ppt and 23-27°C and fed a diet of frozen bloodworms, frozen mysis shrimp, and commercial pellet foods daily. This study used only second-generation through fourth generation lab-reared pupfishes. All newly hatched fry were fed *Artemia* nauplii for approximately one month. All SSI species can be readily crossed to produce viable and fertile hybrids (Martin et al., 2017; St. John et al., 2021). F1 hybrid and F2 hybrid intercrosses were generated from molluscivore x scale-eater crosses from both Osprey Lake and Crescent Pond, generalist x scale-eater and generalist x molluscivore crosses from Little Lake, and generalist x *C*. sp. ‘broadmouth’ F1 hybrids from Osprey Lake (Richards and Martin, 2022). Prior to filming, pupfishes were fed exclusively Repashy Superfood gel diets for acclimation and training for at least one week before filming.

### High-speed filming and measurement of gelatin bites

We recorded pupfishes feeding on standardized gelatin cubes (dimensions: 1.5 cm x 1.5cm x 1.5 cm x 1.5 cm cube; Community Plus Omnivore Gel Premix, Repashy Specialty Pet Products). Gels were prepared in batches of 50 at precisely a 4:1 water:mix ratio in silicone molds following the manufacturer’s instructions and allowed to set overnight at 4° C. Gels were stored covered at 4° C for a maximum of two weeks before discarding. The gel cube retains its shape in water and therefore allows precise measurements of the dimensions and area removed by each bite.

Individuals were trained in group tanks to feed on gelatin cubes and then netted individually for filming. We filmed all strikes at 1,100 fps using a full-color Phantom VEO 440S (Vision Research Inc.) with a Canon EF-S 60 mm lens mounted on a standard tripod. Fish were filmed individually in 7.5 liter bare glass tanks with a solid matte background at salinities of 2-3 ppt and 21-23°C. Illumination was provided by two dimmable full spectrum LED lights placed on either side of the filming tank. Gelatin cubes were held with forceps with one edge facing in a horizontal direction toward the fish (Fig. 1). Trained fish usually attacked the gel almost immediately after placement in the filming tank. After each strike the cube was immediately removed and inspected to confirm a bite mark; missed strikes were confirmed from the video replay. New cubes were used for each feeding strike and never re-used. All videos were filmed in lateral view.

**Fig. 1.**
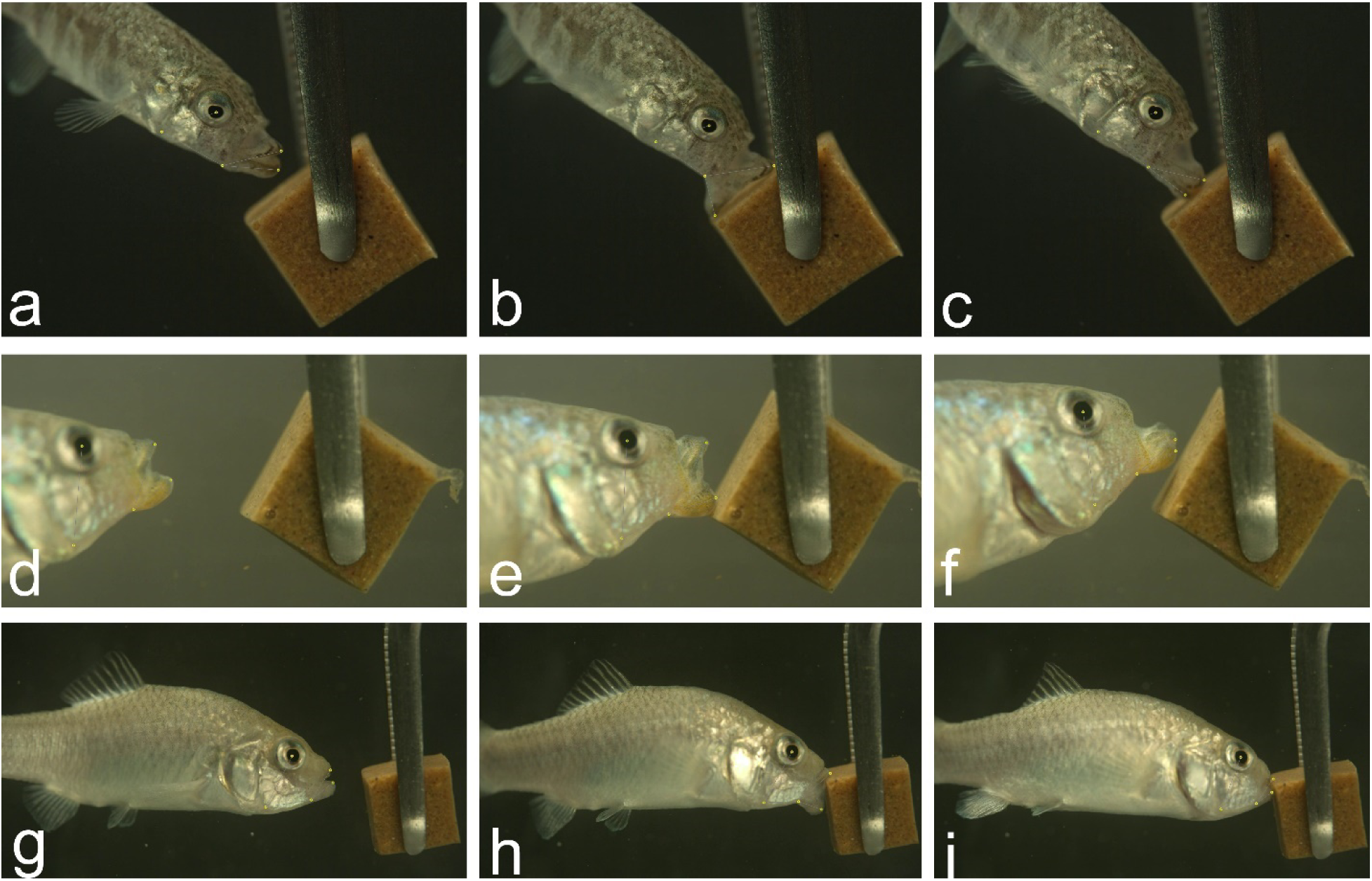
High-speed video of gel-biting strikes with automated landmarks. Strike sequences for a scraping bite by a scale-eater (a-c), a miss by a hybrid (d-f), and an edge bite by a molluscivore (g-i). Videos were filmed at 1,100 fps on a Phantom VEO 440S camera. Frames illustrate approximately 20% of peak gape (first column), peak gape (second column), and jaw adduction immediately after the bite (third column). Five yellow landmarks on each frame were placed automatically using our SLEAP inference model and are illustrated as small yellow dots to emphasize the accuracy of these inferred landmarks.

The length, width, and depth of each gelatin bite were measured using digital high-precision calipers (Mitutoyo) under a stereomicroscope for nearly all strikes (Fig. S1). Strikes were characterized as an edge, corner, scrape, or miss. Edge bites occurred along the edge but not the corner of the gelatin cubes and comprised the majority of strikes. Corner bites were removed from one of the corners, which may affect bite dimensions, so were distinguished from edge strikes. Scrapes were defined by bites in which the jaws did not completely close around the gel to remove a chunk of the material, but instead left two distinct indentations. Misses were defined as strikes in which the oral jaws of the fish contacted the gel but did not leave any marks. Strikes in which the jaws did not make visible contact with the gel were excluded. Most individuals were filmed consecutively over one or two filming periods for up to sixteen strikes. After each filming session for each individual, an image of a ruler was photographed in the filming tank at the same distance as the gelatin cubes for calibration of videos.

### Machine learning for quantifying kinematic landmarks on each frame

We used the SLEAP (Social LEAP Estimates Animal Poses) analysis pipeline to automatically detect and place morphometric landmarks on each frame (Pereira et al., 2020; Pereira et al., 2022). This software supports data input of raw videos and then provides an interactive GUI to create a labeled training dataset. Predictions from trained models can then be adjusted to enable a ‘human-in-the-loop’ workflow to efficiently and progressively obtain more accurate models and inferences of landmarks.

We manually placed five landmarks on 815 frames using the SLEAP GUI spanning 100 high-speed feeding videos including all three species and their hybrids. Frames were chosen for labeling both by eye and automatically by the software to span highly divergent scenes spread across the beginning, middle, and end of each strike. To train a model based on the labeled data, after exploring various configurations we achieved the best performance using the multi-animal bottom-up unet model with a receptive field of 156 pixels, max stride of 32 pixels, batch size of 3, input scaling of 0.75, and validation fraction of 0.1, resulting in a precision of 99% and mean distance between labeled and inferred landmarks of 5 pixels (Fig. S2). We trained this model on a laptop running a 16 Gb NVIDIA GeForce 3070 GPU, which completed training in less than twelve hours. We then used this trained model to infer landmarks on each frame of a larger set of strike videos using the flow cross-frame identity tracker, which shares information about landmark places across frames for each individual strike video. We culled to a single instance (i.e. one animal) per frame, given that fish were filmed individually, and connected single track breaks. We predicted landmarks on 114,000 frames from 227 .mp4 videos (batch converted from the original .cine files using Phantom Camera Control software) and spot-checked for accuracy (Fig. 1). Coordinate data from each frame were exported in .hdf5 format, imported into R (Core Team, 2021), and stored in any array using the rhdf5 package (Fischer et al., 2017).

### Kinematic variables

To quantify gel-biting strikes from coordinate data, we calculated five key kinematic variables per strike: 1) peak gape, the distance from the anterior tip of the premaxilla to the anterior tip of the dentary; 2) peak jaw protrusion, the distance from the center of the pupil to the anterior tip of the premaxilla; 3) peak lower jaw angle, the minimum angle between the lower jaw, the quadrate-articular point of jaw rotation, and the ventral surface of the fish beneath the suspensorium measured from an anteroventral landmark on the preopercle. This measures the maximum rotation of the oral jaws in a downward and outward direction toward the gel as defined in St. John et al. (2020). Note that in Cyprinodontiformes oral jaw opening is decoupled from jaw protrusion by the maxillomandibular ligament such that peak gape does not necessarily occur simultaneously with peak jaw protrusion (Hernandez et al., 2009). 4) Time to peak gape (TTPG) was the time in ms from 20% of peak gape to peak gape. 5) Ram speed (m/s) was calculated as the distance from 20% of peak gape to peak gape (mm) divided by the time to peak gape (ms). We calibrated each set of coordinates for each filming session using a ruler held at the same distance from the camera as the gelatin cube. Milliseconds were calculated by counting frames and multiplying by 0.909 to correct for the 1,100 frame rate per second. We then subsetted to only those frames from initial start position to maximum distance from the start position before the fish started to turn its head to the side post-bite so that kinematic variables were only calculated from the start of the strike to the time of impact with the gelatin cube (see Supplemental R code).

## Statistical analyses

### Mixed-effect modeling

Due to our repeated measures design of multiple strikes per fish, we used mixed-effects models to compare kinematic variables and bite dimensions across species groups. We used the lme4 and lmerTest packages in R to fit linear mixed-effects models for each kinematic response variable and bite metric with independent fixed effects for strike type and species (scale-eater, molluscivore, generalist, or hybrid) plus the random intercept effect of individual ID. We measured up to 16 strikes per individual fish. P-values were assessed for each factor level using Satterthwaite’s method (Kuznetsova et al., 2017). We used AIC to compare additional models with random slopes and interactions among the fixed effects (Burnham et al., 2011).

Finally, due to the failure of this radiation to fit a tree-like model of evolution due to extensive secondary gene flow (Richards et al., 2021), in addition to our inclusion of several hybrid crosses, we did not correct for phylogenetic signal in our analyses. However, we note that both generalist outgroup populations are more distantly related to each other than the scale-eater, molluscivores, and hybrids on SSI (Martin and Feinstein, 2014), indicating that kinematic variables and bite dimensions exhibit minimal phylogenetic signal (Losos, 2008).

### Multivariate analyses of kinematic variation

We calculated principal component analysis (princomp) from the correlation matrix of kinematic variables. We also used linear discriminant analysis from the MASS package in R (Ripley et al., 2013) to explore overall kinematic variation among species and strike type. We further calculated classification accuracy using species or strike type as the grouping variable using all five kinematic variables.

### Generalized additive modeling

We used generalized additive semi-parametric models (GAMs) to test for nonlinear terms relationships and interactions between kinematic variables and bite dimensions (Wood and Augustin, 2002). Because we were interested in directly predicting bite performance from the kinematic data for each strike, we treated strike as our unit of replication, not fish, and did not control for individual in our statistical models. However, within-individual variation often exceeded between-species variation and we were not directly interested in species kinematic differences using this modeling framework, which we previously addressed explicitly using mixed-effects models controlling for individual. Moreover, methods for fitting mixed-effects GAM models in R are currently limited.

We fit GAM models using the mgcv package in R (Wood, 2012; Wood and Wood, 2015) with the response variables of bite length, width, depth, or volume and independent covariates of species and strike type, and independent continuous kinematic variables of peak gape, peak jaw protrusion, peak lower jaw angle, time to peak gape, and ram speed. We used the REML method for calculating smoothness of splines and Gaussian distributions for all models. We compared the fit of each kinematic variable modeled as both a linear term or a smoothing spline using model selection with the AIC criterion in R. We further allowed for shrinkage of each smoothing spline within the full model to determine which kinematic variables were best modeled as spline terms. We then systematically compared models with both two-way thin-plate splines or all one-way splines to explore whether there were any nonlinear interactions between kinematic variables. Model fits were visualized with the mgViz (Fasiolo et al., 2020) and ggplot2 packages (Wickham et al., 2016) in R. All coding was assisted by suggestions from ChatGPT 3.5 and 4.0 (OpenAI, Inc.).

## Results

### Scale-eaters displayed increased gape size and bite length

In mixed-effects models controlling for species, strike type, and repeated sampling of each individual, we found no effect of species on bite depth or width (effect of scale-eater factor level: *P* = 0.062); however, there was a significant positive effect of scale-eater species identity on bite length (*P* = 0.0060). Among the five kinematic variables, only increased peak gape was significantly associated with scale-eater strikes (*P* = 0.040). No kinematic variables were significantly associated with any other species or hybrids across strike types (Fig. 2), even when comparing missed strikes to successful bites.

**Fig. 2.**
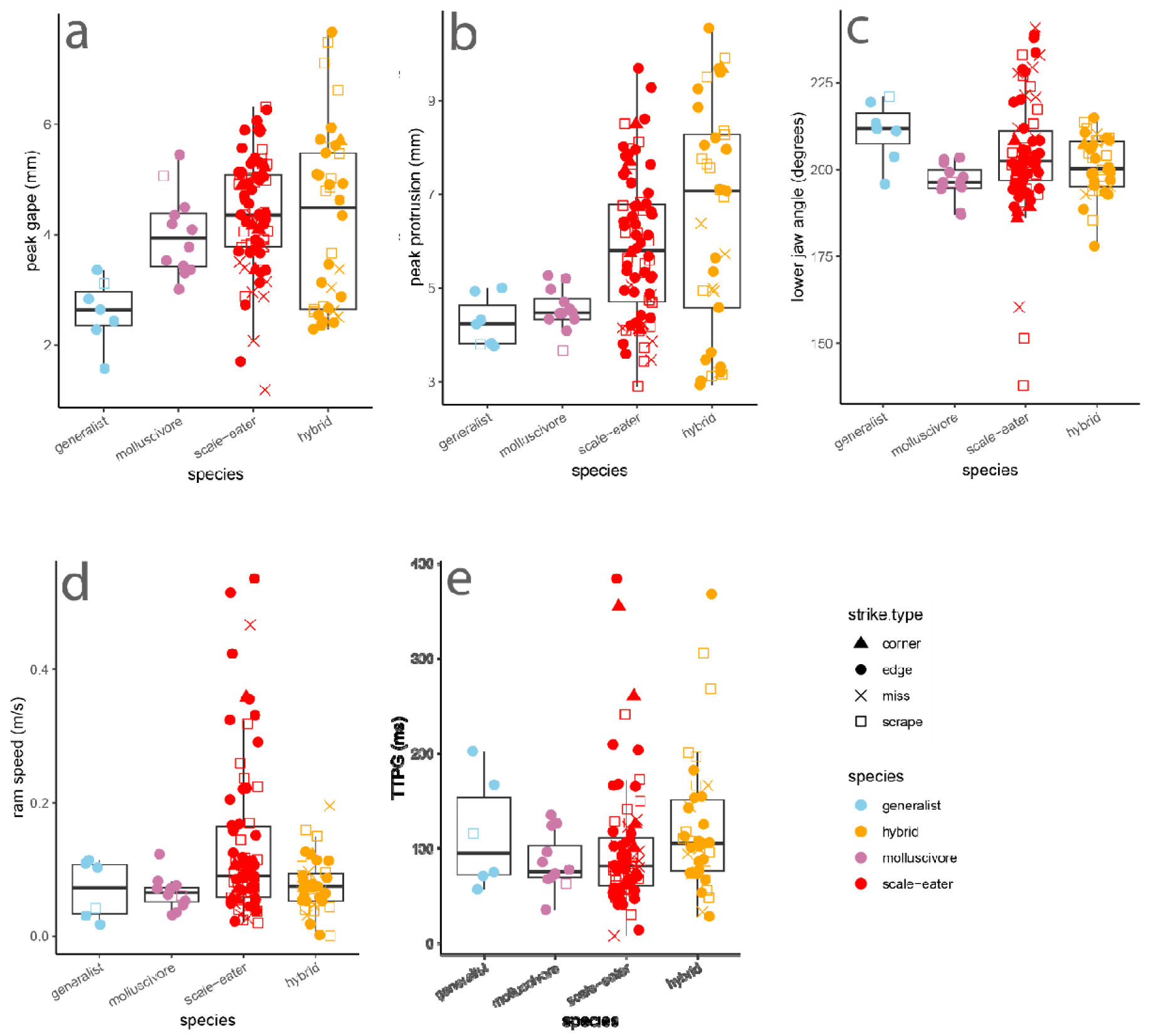
Scale-eaters only differ in increased peak gape during gel-biting strikes. Boxplot overlaid with raw data show five kinematic variables measured during gel-biting strike measured from automated landmarking of 227 videos. Species or hybrid cross is indicated by color and strike type is indicated by shape. TTPG: time to peak gape. Lower jaw angle is the minimum angle between the lower jaw and suspensorium from 20% to peak gape. Ram speed was calculated from the distance traveled between 20% and peak gape.

### Substantial similarity in strike kinematics among species and strike types

Similarly, we found substantial overlap in kinematic variation across species and strike types. Principal component analysis showed the strongest loadings of peak gape and peak jaw protrusion on PC1, explaining 36.7% of kinematic variance among strikes (Fig. 3).

**Fig. 3.**
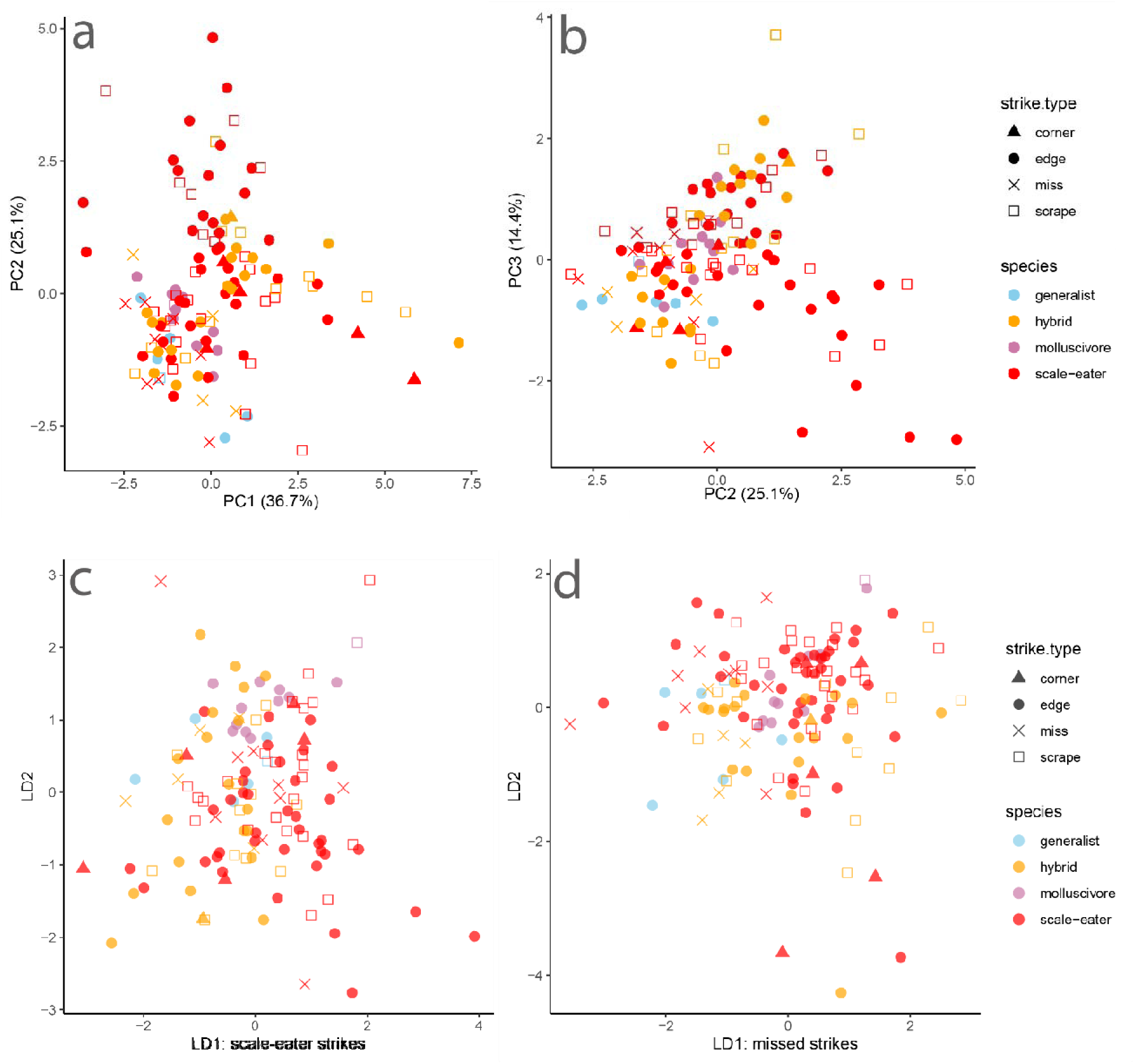
Principal component and linear discriminate analyses illustrate substantial overlap by strike type and species. Species or hybrid is indicated by color and strike type is indicated by shape. Multivariate analyses were based on five kinematic variables: peak gape, peak jaw protrusion, minimum lower jaw angle with the suspensorium, time to peak gape (TTPG), and ram speed.

Linear discriminant analysis by strike type successfully classified strikes at a rate of only 50.8%. The kinematic variable best separating misses from other strike types on discriminant axis one was peak gape. Linear discriminant analysis by species successfully classified species or hybrids based on their strike kinematics at a rate of only 32.3%. The kinematic variables best separating scale-eaters from other groups on discriminant axis one was again peak gape and peak jaw protrusion while TTPG had the weakest effect on classification of species by kinematic variables. Although plots of the first two principal components and discriminant axes indicate greater variation within scale-eater and hybrid strike kinematics, there was also clearly substantial overlap among species and hybrids (Fig. 3).

### Multi-peak performance landscape for gel-biting

We used generalized additive modeling to explore the relationship between kinematic variables and bite size dimensions. The best fit model (Table 1) included a two-dimensional thin-plate spline for peak gape and peak jaw protrusion along with fixed linear predictors for species, strike type, peak lower jaw angle, time to peak gape (TTPG), and ram speed. The nonlinear interaction between peak gape and peak jaw protrusion was significantly associated with both the bite length (edf = 10.82, *P* = 9e^-7^) and the overall gel volume removed (edf = 8, *P* = 0.0008) and displayed a bimodal surface with two isolated performance peaks (Fig. 4). The best fit model for bite length included additional significant linear effects of ram speed (*P* = 0.008) and peak lower jaw angle (*P* = 0.007), but not TTPG (*P* = 0.440) in addition to significant factor levels of missed strikes (*P* = 1.12e^-10^) and molluscivore species (*P* = 0.012). Models without TTPG fit the data equally well (ΔAIC < 2).

**Fig. 4.**
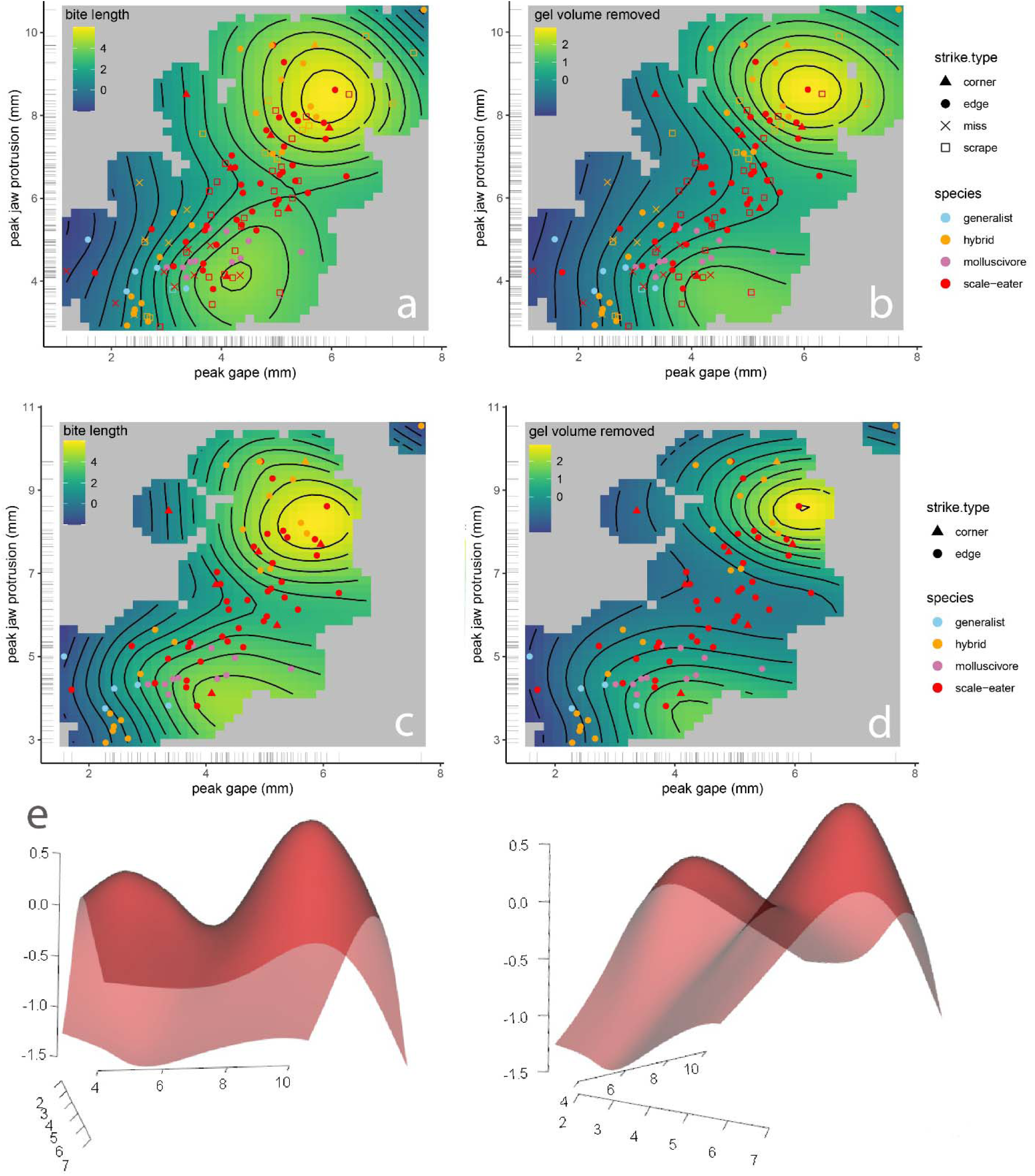
Generalized additive modeling supports a nonlinear interaction between peak gape and peak protrusion for bite performance. Thin-plate splines from the best-fitting GAM model (Table 1) for the response variable of bite length (first column: **a,c**) or total bite size (gel volume removed: second column: **b,d**). First row includes all strike types and second row excludes scraping and missed strikes. **e.** Performance landscape for bite length in 3D perspective view. Species or hybrid cross is indicated by color and strike type is indicated by shape for 130 filmed strikes with data for gel bite dimensions. Gel volume removed was calculated from length x width x depth of gelatin bite measured with a digital caliper under a stereomicroscope.

**Table 1.**
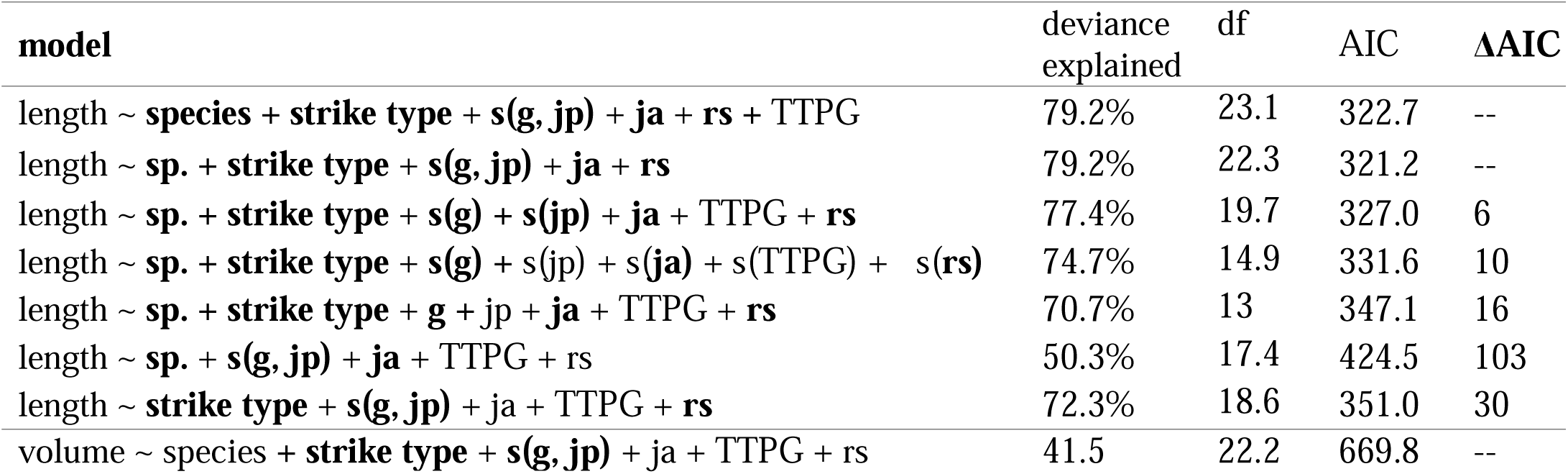
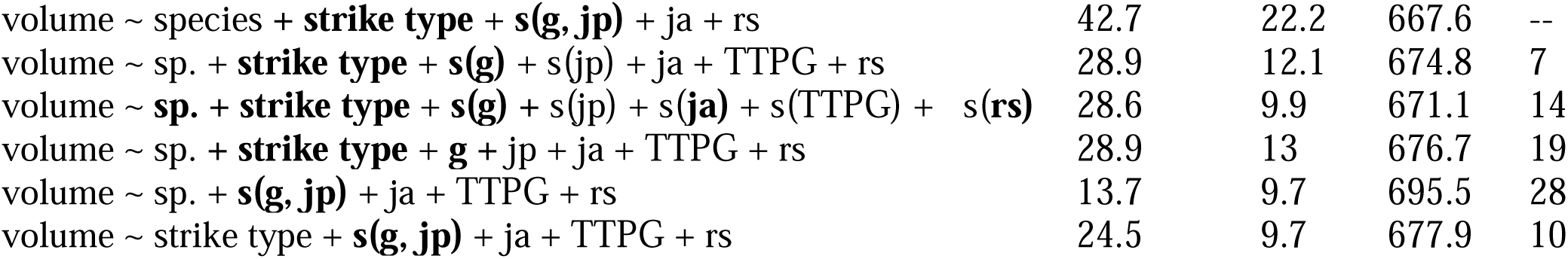
Model selection of generalized additive models (*n* = 130 strikes with complete kinematic and gel bite data) predicting bite length or total bite volume removed from Repashy gelatin cubes by a single strike. sp: species; g: peak gape, jp: peak jaw protrusion; ja: peak lower jaw angle with the suspensorium; TTPG: time to peak gape; rs: ram speed. Strike type included full bites from the gelatin edge, corner, scraping bites, and misses in which the jaws made contact with the gelatin but left no impression. Significant terms within each GAM model (*P* < 0.05) are highlighted in bold.

| model | deviance explained | df | AIC | $\Delta AIC$ |
| --- | --- | --- | --- | --- |
| length ~ **species** + **strike type** + **s(g, jp)** + **ja** + **rs** + TTPG | 79.2% | 23.1 | 322.7 | -- |
| length ~ **sp.** + **strike type** + **s(g, jp)** + **ja** + **rs** | 79.2% | 22.3 | 321.2 | -- |
| length ~ **sp.** + **strike type** + **s(g)** + **s(jp)** + **ja** + TTPG + **rs** | 77.4% | 19.7 | 327.0 | 6 |
| length ~ **sp.** + **strike type** + **s(g)** + **s(jp)** + **s(ja)** + **s(TTPG)** + **s(rs)** | 74.7% | 14.9 | 331.6 | 10 |
| length ~ **sp.** + **strike type** + **g** + **jp** + **ja** + TTPG + **rs** | 70.7% | 13 | 347.1 | 16 |
| length ~ **sp.** + **s(g, jp)** + **ja** + TTPG + **rs** | 50.3% | 17.4 | 424.5 | 103 |
| length ~ **strike type** + **s(g, jp)** + **ja** + TTPG + **rs** | 72.3% | 18.6 | 351.0 | 30 |
| volume ~ species + **strike type** + **s(g, jp)** + **ja** + TTPG + rs | 41.5 | 22.2 | 669.8 | -- |
| volume ~ species + **strike type** + **s(g, jp)** + ja + rs | 42.7 | 22.2 | 667.6 | -- |
| volume ~ sp. + **strike type** + **s(g)** + s(jp) + ja + TTPG + rs | 28.9 | 12.1 | 674.8 | 7 |
| volume ~ **sp.** + **strike type** + **s(g)** + s(jp) + s(**ja**) + s(TTPG) + s(**rs**) | 28.6 | 9.9 | 671.1 | 14 |
| volume ~ sp. + **strike type** + **g** + jp + ja + TTPG + rs | 28.9 | 13 | 676.7 | 19 |
| volume ~ sp. + **s(g, jp)** + ja + TTPG + rs | 13.7 | 9.7 | 695.5 | 28 |
| volume ~ strike type + **s(g, jp)** + ja + TTPG + rs | 24.5 | 9.7 | 677.9 | 10 |

Even after excluding missed strikes that made contact but left no mark on the gel and scraping bites in which the jaws did not fully occlude, the interaction between peak gape and jaw protrusion was still significantly associated with edge and corner bite length (edf = 10.34, *P* = 9e^-6^) and volume (edf = 11.17, *P* = 4.89e^-5^). Bite width was significantly linearly associated with the interaction between peak gape and peak jaw protrusion resulting in a flat performance landscape (edf = 1.759, *P* = 0.0002), which is largely controlled by the morphological dimensions of the oral jaws of each fish, rather than kinematic variables. Bite depth was not significantly associated with peak gape and peak jaw protrusion nor any linear variable in this model except for the factor of scraping strikes as expected based on the shallow dimensions of these strikes (*P* = 0.046).

## Discussion

We estimated a surprisingly complex performance landscape for the unusual trophic niche of lepidophagy from high-speed videos of gelatin-removing bites. In contrast to studies of suction-feeding performance on evasive, attached, and strain-sensitive (e.g. zooplankton) prey (Holzman et al., 2008; Holzman et al., 2022; Keren et al., 2018; Olsson et al., 2020), we found no effect of kinematic timing variables such as TTPG or even ram speed on the performance of scale-biting strikes, measured by the length, width, depth, and total volume of gelatin removed. Instead, successful scale-biting appears to require strike coordination between jaw opening (peak gape) and jaw protrusion and, surprisingly, this interaction resulted in two distinct performance optima: 1) individuals with small peak gapes removed the greatest amount of material per bite at small jaw protrusion distances; 2) individuals with large peak gapes removed the greatest volumes at large jaw protrusion distances; and 3) these two performance optima were surrounded by reduced bite performance in all directions including more extreme values and intermediate values of peak gape and protrusion. This resulted in two distinct performance peaks on the two-dimensional thin-plate spline for jaw protrusion and peak gape in the model best supported by the data (Fig. 4, Table 1). Thus, the strikes with the largest peak gapes and jaw protrusion distances observed suffered a performance decline, in line with a few datapoints from our initial kinematic study of biting in this system (St. John et al., 2020); similarly, the strikes with the smallest peak gapes and jaw protrusion distances also suffered a performance decline. Therefore, we unexpectedly found evidence of a multi-peak performance landscape for the relatively straightforward functional task of biting, a well-studied functional task which is often viewed as a simple mechanical system (Herrel et al., 2008; Wainwright and Richard, 1995; Westneat, 2005).

### Two distinct performance optima for biting rather than a linear ridge

Surprisingly, there was no simple linear ridge for the interaction between peak gape and peak jaw protrusion in relation to bite volume or bite length; both extreme and intermediate values of these kinematic variables resulted in poor performance, i.e. reduced gelatin bite sizes (Fig. 4). Only strikes by scale-eating specialists and some hybrid strikes resided on the second performance peak with larger gapes and jaw protrusion while all generalists and molluscivores occurred on or near the first performance peak. This suggests that a performance valley isolates the recently evolved scale-eating specialist *C. desquamator* from its generalist ancestor. Explanations for this performance landscape must also account for the poor performance of intermediate strike values observed, rather than just a simple performance ridge indicating that coordination between peak gape and peak jaw protrusion is important.

One possible explanation is a biomechanical tradeoff in precision and targeted bite area with the most adverse effects on gel-biting performance at intermediate values. Interactions between oral jaw scraping and biting with the gelatin surface may only be effective within two different kinematic regimes. Smaller peak gapes with less jaw protrusion may allow for precise targeting and higher mechanical advantage for removing more gelatin. Larger peak gapes with greater jaw protrusion may reduce precision and mechanical advantage of the bite, but cover a large area, resulting in more gelatin removed per bite. Intermediate values may suffer the costs of less precise biting and less area covered per bite. It is tempting to speculate that strike speed or lower jaw angle play a role in this precision/target area tradeoff. However, while ram speed and lower jaw angle with the suspensorium both had strong linear effects on bite performance, there was no evidence of any nonlinear interactions with peak gape or peak jaw protrusion distance (Table 1). Similarly, timing (TTPG) seems to play no role in bite performance, which would seem surprising if precision is important for gelatin removal since faster time to peak gape should reduce bite precision. However, biting strikes generally achieved peak gape substantially before contact with the gel, suggesting that the time to reach peak gape does not seem to be important as long as the jaw is open at some point before contact with the target. This is the consistent with the ‘plateau effect’ observed in the scale-eating piranha, the only other direct study of scale-biting kinematics in other systems (Janovetz, 2005).

Alternatively, it is striking that none of the generalist or molluscivore species exhibited feeding strikes with jaw protrusion distances within range of the second performance peak (Fig. 4). Similarly, only hybrids with scale-eater ancestry were capable of producing feeding strikes with jaw protrusion distances in this range (> 6.5 mm; see supplemental raw data). Thus, there appears to be a genetic basis underlying the two performance peaks: only scale-eaters and hybrids with scale-eater ancestry can protrude their jaws sufficiently to reach the second performance optimum. This may be due to additional anatomical properties of their oral jaws that allow for greater extension during strikes, such as different ratios of muscle fiber types within the adductor mandibulae (Ono and Kaufman, 1983; Summers and Long, 2005), along with unmeasured aspects of their behavior or strike kinematics. Indeed, scale-eaters exhibit significant differences in their boldness and exploratory behaviors (St. John et al., 2020). Genome-wide association scans for oral jaw length also identified collagen genes with fixed regulatory differences between scale-eaters and molluscivores, including collagen type XV alpha 1 (col15a1), suggesting that the elasticity of jaw opening may be under selection in this species (Martin et al., 2019; McGirr and Martin, 2017). Greater peak gapes are possible due to the two-fold larger oral jaws of the scale-eater. However, scale-eaters still do not open their jaws as wide as possible during strikes (St. John et al., 2020) or achieve 180° angles with their open jaws as in other scale-eating specialists such as the scale-eating piranha (Janovetz, 2005), indicating adaptive behavioral compensation for their extreme oral anatomy during strikes.

Finally, we cannot rule out more esoteric explanations for the unexpected fitness valley between bite performance optima. Sensory perception during strikes may be limited at intermediate strike distances due to the blind spot caused by the positioning of the vertebrate optic nerve in front of the retina, although biomechanical implications of this in fishes are unknown (Gregory and Cavanagh, 2011). Alternatively, intermediate jaw protrusion may be an indirect effect of premature suspension of strike behavior or lack of motivation during the strike. However, excluding missed strikes did not alter the observed two-peak performance landscape (Fig. 4).

### Similarity between the performance and fitness landscapes

Both field measurements of fitness and laboratory measurements of scale-biting performance support a two-peak landscape. Repeated field experiments in this system measured the fitness landscape from the growth and survival of advanced generation hybrids placed within 3-4 m enclosures in their natural hypersaline lake environments for 3 – 11 months and estimated two fitness peaks separated by a fitness valley (Martin and Wainwright, 2013b). Surprisingly, these landscapes remained relatively stable and exhibited a similar two-peak topography across years, lakes, and frequency-manipulations of hybrids (Martin, 2016a; Martin and Gould, 2020). However, in all cases where it was detected, the second peak corresponded to the phenotype of molluscivores; whereas hybrids resembling the scale-eater survived and grew at the lowest rates across all fitness experiments. Thus, a single fitness peak corresponds to generalist morphology and kinematics for both fitness and performance, whereas the second scale-biting performance peak has no analog in field enclosures, perhaps due to a lack of hybrids with the necessary combination of scale-biting kinematics and morphology. However, these laboratory estimates do suggest that F2 hybrids with scale-eater ancestry display sufficient jaw protrusion distances during gel-biting strikes to occupy the second performance peak.

Ultimately, the goal of connecting morphological fitness landscapes to performance landscapes is difficult due to the many-to-one mapping of hybrid morphologies onto biting kinematics. However, better understanding of the genetic basis of kinematic variables, such as jaw protrusion distance, may enable connecting the genetic regulatory networks underlying morphological, kinematic, and behavioral traits through genotypic fitness networks informed by laboratory performance and field fitness experiments. Our initial study of genotypic fitness networks found rare but accessible pathways in genotype space (i.e. equal or increasing in fitness at each step) connecting scale-eaters to other species, but so far we have found no associations between fitness and any behavioral genes (Patton et al., 2022). Also see the role of *sox9b* in cichlid foraging performance, in which correcting for genotype improves the form-function relationship (Matthews et al., 2023).

## Conclusion

Here we explore the biomechanics of a highly specialized trophic niche and demonstrate the power of machine-learning approaches to analyze kinematic data. We estimated a surprisingly complex two-peak performance landscape for biting that indicates that the highly protrusible jaws of scale-eating specialists may provide a performance benefit for scale-eating. This study provides a new framework for understanding bite mechanics in fishes – particularly scraping dynamics – and a foundation for dissecting the genetic basis of these predatory behaviors and their relationship to fitness landscapes driving rapid adaptive radiation in the wild.

## Acknowledgments

We thank Jack Tseng and the Martin and Holzman labs for valuable comments and discussion of the results. We also thank the Gerace Research Centre and Troy Day for logistical support and the government of the Bahamas for permission to collect and export breeding colonies in 2017 and 2018. This research was funded by the Binational Science Foundation 2016136 to RH and CHM, and the National Science Foundation DEB CAREER grant #1749764, National Institutes of Health grant 5R01DE027052-02, the University of North Carolina at Chapel Hill, and the University of California, Berkley to CHM and a Discovery for All grant for discovery-based learning to AT.

---

**Fig. S1.**
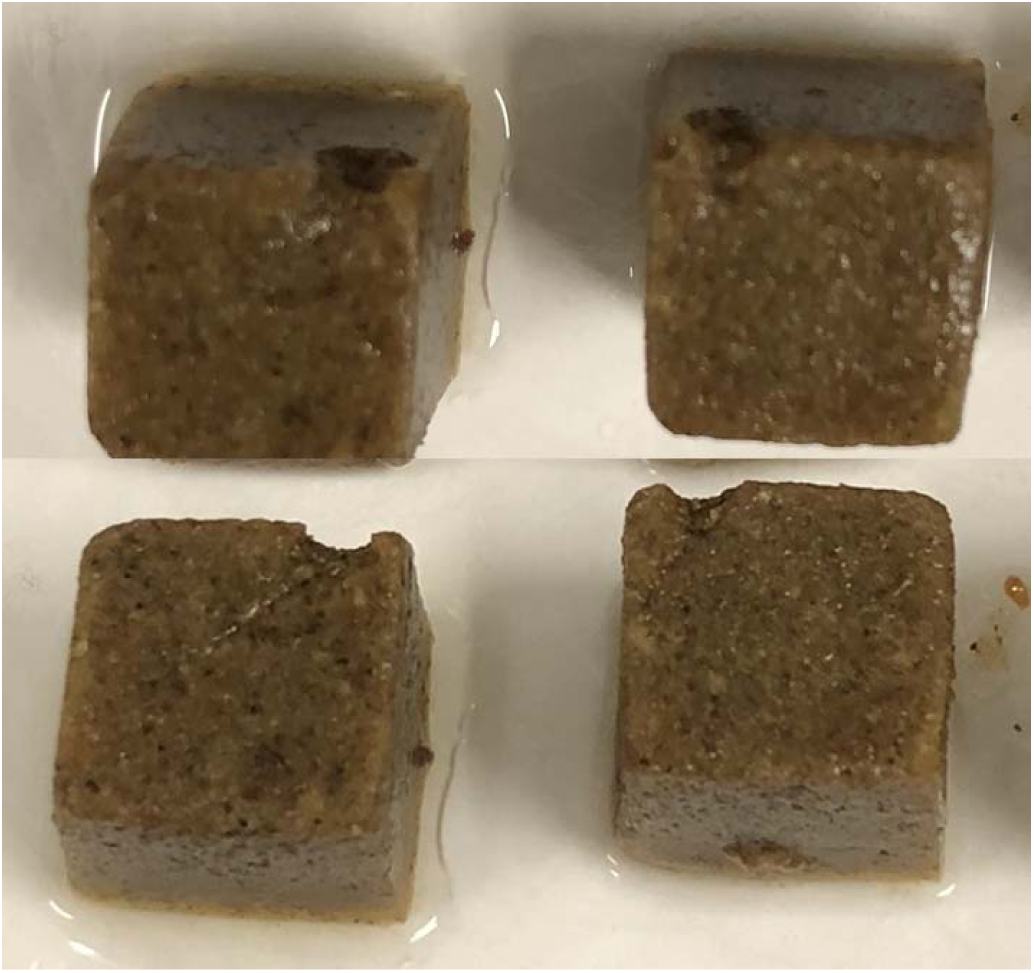
Top-down and side views of typical single bites from Repashy gelatin cubes. Both bites pictured were classified as edge bites, the most common type of bite recorded.

**Fig. S2.**
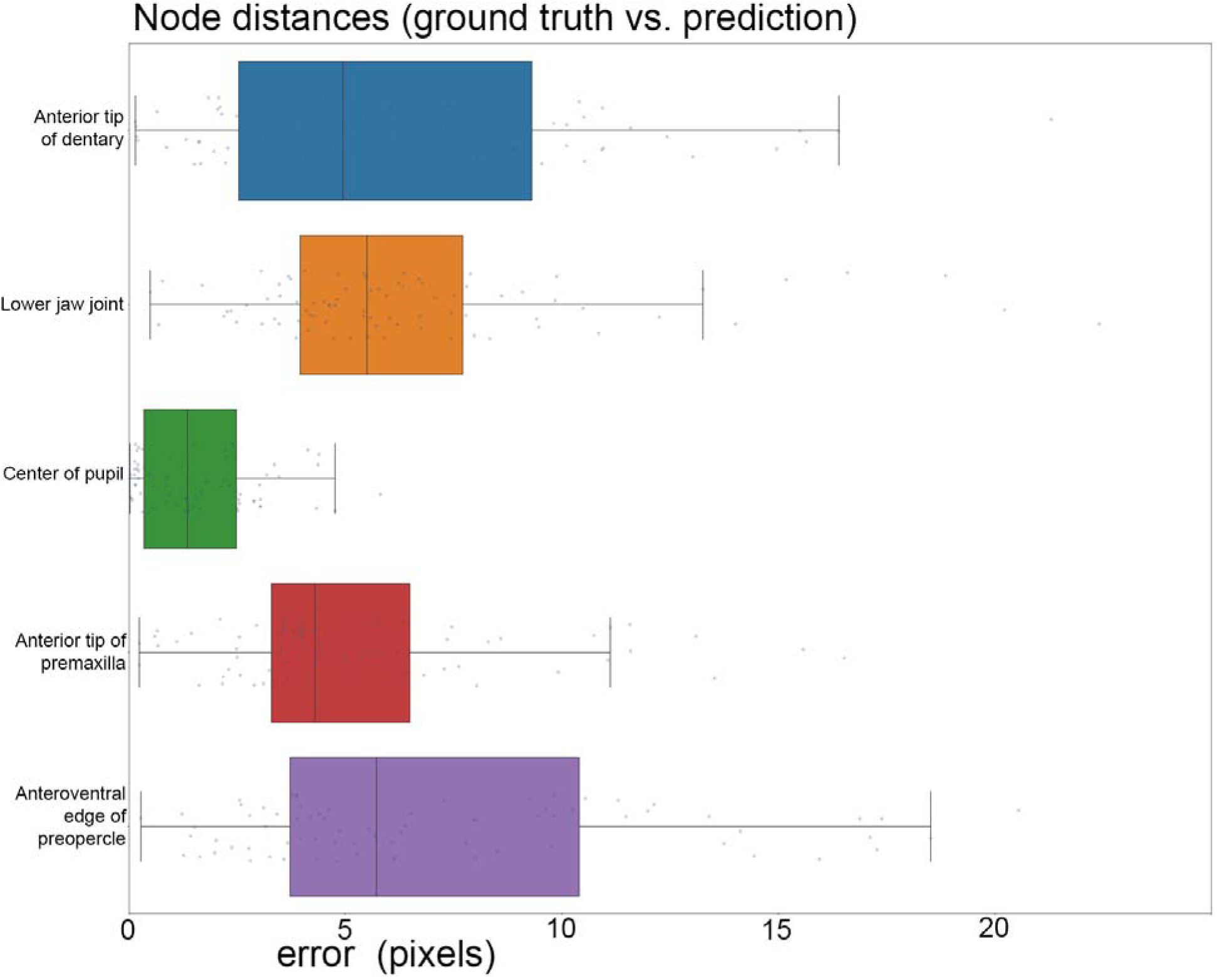
Error distribution for each landmark in our best performing model. Trained model was based on 815 labeled frames across 100 high-speed videos. We used the multi-animal bottom-up unet model with a receptive field of 156 pixels, max stride of 32 pixels, batch size of 3, input scaling of 0.75, and validation fraction of 0.1. The mean distance between labeled data and inferred landmark positions was 5.80 pixels. Precision obtained was 0.990 based on four false positives, five false negatives, and 397 true positives in the model summary output from SLEAP.

